# New insights into the dynamics of *de novo* gene origin

**DOI:** 10.1101/2023.12.08.570739

**Authors:** Logan Blair, Julie Cridland, Yige Luo, David Begun, Artyom Kopp

## Abstract

The evolution of genes *de novo* from ancestrally nongenic sequences is a significant mechanism of gene origin. Many studies have focused on distant evolutionary comparisons, which bias the sample of *de novo* genes towards older genes that have acquired important functions and have been refined by selection. In this report, we focus on the earliest steps in *de novo* gene origin by identifying young, polymorphic transcripts that may be missed by other study designs. To accomplish this, we sequenced tissue-specific transcriptomes from a much larger sample of genotypes than have been used in previous analyses of *de novo* genes in *Drosophila melanogaster*. We identified 90 potential species-specific *de novo* genes expressed in the male accessory glands of 29 *D melanogaster* lines derived from the same natural population. We find that most young, unannotated transcripts are both rare in the population and transcribed at low abundance. Improved sampling of both ingroup and outgroup genotypes reveals that many young genes are polymorphic in more than one species, resulting in substantial uncertainty about the age and phylogenetic distribution of *de novo* genes. Among the genes expressed in the same tissue, gene age correlates with proximity to other tissue-specific genes, with the youngest genes being least likely to occur near established tissue-specific genes. This and other lines of evidence suggest that *de novo* genes do not commonly evolve by simply reutilizing pre-existing regulatory elements. Together, these results provide new insights into the origin and early evolution of *de novo* genes.

**Author Summary:** Genes may be born and lost without any lasting evidence of their existence. The typical longevity may be especially limited for *de novo* genes – that is, genes that originate from ancestrally non-genic, untranscribed sequences, since most genomic regions are not expected to be beneficial when transcribed. To better capture the population biology of nascent *de novo* genes at points close to their origin, we sequenced tissue-specific transcriptomes from a large number of *Drosophila melanogaster* genotypes. Most *de novo* genes were expressed in very few genotypes, consistent with the expectation of transience and rapid turnover. However, many young genes showed polymorphic transcription in multiple species, suggesting that the combination of low frequency with limited sampling can lead us to underestimate how long *de novo* genes persist in populations. We identified several features that *de novo* genes come to share with established tissue-specific genes the longer they persist. This study highlights important challenges in reconstructing *de novo* gene origin and helps elucidate why some transcripts may survive long enough to acquire selectable functions.

## Introduction

Widespread genome and transcriptome sequencing has shown that new genes can originate from DNA sequences that, in the ancestral state, show no evidence of transcription (1). The evolution of these “*de novo* genes” from nongenic sequences has now been documented across the tree of life (2–5). However, questions remain about how prevalent *de novo* origin is compared to the long-accepted evolution of new genes by duplication (6), how much these genes contribute to organismal fitness (7), and how they originate.

Though *de novo* genes with crucial function have been identified (8,9), less is known about their initial origin. Partly this is due to study design. Many studies have focused on *de novo* genes where the most recent common ancestor is more than a few million years old, including studies within model organisms such as humans (10,11) and mice (12,13). These timescales may afford greater time for the *de novo* gene sequences to be refined by selection, and thus a greater proportion of *de novo* genes identified may be of phenotypic importance. However, these studies have drawbacks for elucidating the early stages of gene origin. Long divergence between lineages with and without the *de novo* genes allows extensive sequence differences to accumulate, obscuring the earliest mutations that spawned the novel transcripts. Greater divergence also makes it increasingly difficult to determine orthology between new genes and their progenitor nongenic sequences and may increase the rate of false positive *de novo* gene calls. Additionally, some studies that do incorporate more recent outgroups do not include noncoding RNAs (14,15). Functional proteins are only a subset of all *de novo* genes, and such limited focus may lead to a biased picture of the evolutionary trajectories that *de novo* genes can follow. For instance, some *de novo* genes may become transcribed first and acquire ORFs later (16,17). With these points in mind, there is a need for studies on shorter evolutionary timescales, particularly those that incorporate rare, low-abundance, and noncoding transcripts.

Since the youngest *de novo* genes are expected to be expressed in a small proportion of individuals of a single species, studies aimed at understanding the earliest steps in gene origin must be designed to capture such genes. This poses both conceptual and technical challenges because the presence or absence of a gene in a given species cannot be approached as a binary trait. The prevalence of rare *de novo* gene alleles (5,18) suggests that most of them are transient and will eventually be lost. Those that fix in all individuals of a species necessarily go through a polymorphic stage. Sampling a young *de novo* gene is therefore probabilistic. At the same time, the absence of an orthologous transcript in related species must be regarded as a working hypothesis that may be rejected by improved sampling. Finally, if an ancestral polymorphic *de novo* gene segregates in two sibling species, with increasing divergence time, there is a growing chance that this gene will be lost in one of the lineages. Once this occurs, the *de novo* gene in the other species will appear to be significantly younger than it really is.

The population biology of young *de novo* genes has previously been studied in *Drosophila melanogaster*, which serves as an excellent model due in part to the availability of many closely related species with well annotated genomes. While these studies have examined a variety of tissues, including whole-body (14) and female reproductive tracts (19), particular focus has been given to the rapidly evolving male reproductive tissues of the testis (5) and the accessory gland (AG) (18), which produces seminal fluid (20). Despite the involvement of both the testis and the AG in male reproduction, the population biology of *de novo* genes appears to differ between the two organs. The testis expresses more *de novo* genes, those genes are present at higher frequencies, and their expression tends to have a stronger *cis*-regulatory component compared to AG-expressed *de novo* genes (5,18). In both tissues, however, many *de novo* genes were transcribed in only one or two of the sampled individuals, suggesting that more polymorphic *de novo* genes are likely to be identified with more extensive sampling.

In addition to the question of “when”, another open question is “how”: are there attributes of ancestral sequences that increase the probability of *de novo* gene origination? Recent studies have suggested that new transcription may be facilitated by preexisting regulatory features such as bidirectional promoters (12,21,22), open chromatin (23), and *cis*-regulatory elements (24). As such, it is possible that *de novo* genes tend to originate near other genes where these regulatory features are more likely to be located (18,25). While studies indicate that reusing existing regulatory machinery may be a path of least resistance in *de novo* gene evolution, the relative importance of different genomic features is not clear. It is also unclear whether these features affect the probability of initial transcription, versus the probability that a *de novo* gene will persist in the population after becoming transcribed. Therefore, there is a need for more documentation of the genomic environment of *de novo* genes over varying evolutionary timescales.

To better understand the early stages of *de novo* gene origin, we sequenced the AG transcriptomes of 29 *D. melanogaster* strains derived from a single natural population. We take a comprehensive, lenient approach to unannotated transcript discovery: we use weak minimum expression filters and do not require *de novo* transcripts to have ORFs. At the same time, we employ strict outgroup filtering procedures with an expanded pool of outgroup sequences, ensuring high stringency for determining whether a polymorphic *de novo* gene is species-specific. With this larger pool of polymorphic transcripts, we explore how species-specific *de novo* genes may differ from other gene classes. We ask: (1) how the probability of sampling unannotated genes depends on their frequency in populations and estimated age, and (2) to what extent various proxies for preexisting regulatory information (such as older tissue-specific genes and bidirectional promoters) increase the likelihood of finding nearby *de novo* genes.

## Results

### Interspecific and intraspecific distribution of unannotated transcripts

We sequenced AG transcriptomes from 34 different *Drosophila* genotypes to better understand how potential *de novo* genes are distributed within and between species. Genotypes within our focal species *D. melanogaster* were sampled most extensively, where we individually sequenced 29 lines from the DGRP population. Using these data, as well as a number of transcriptomes from other studies (Table 1 and S1-S2 tables), we identified unannotated genes specific to the *melanogaster* species subgroup (*D. melanogaster*, *D. simulans*, and *D. yakuba* in this study).

**Table 1:**
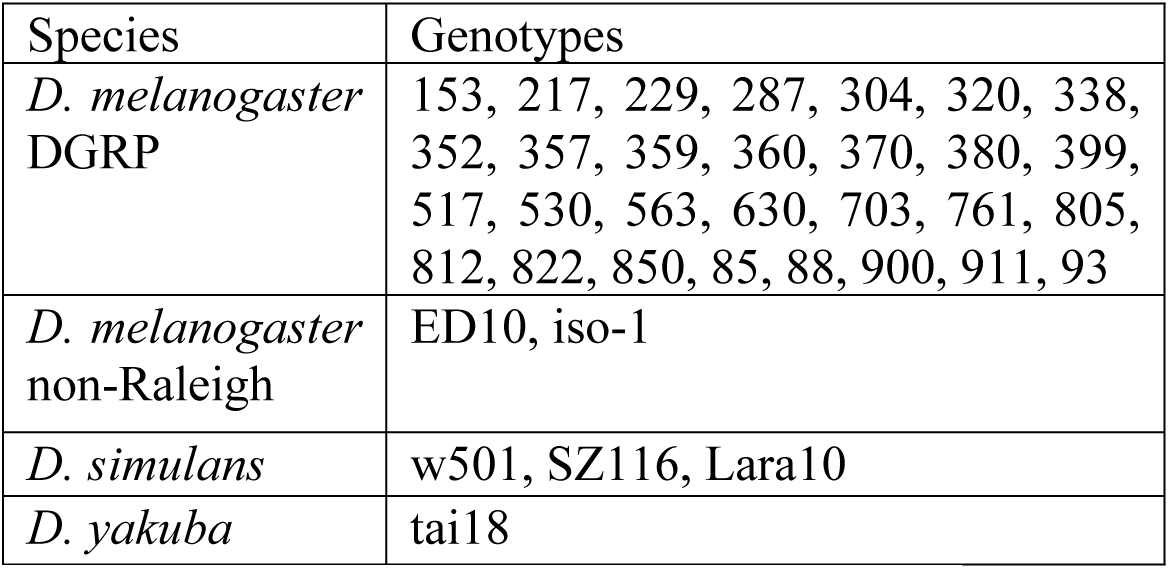
Lines used in this study. DGRP lines are described in Mackay 2012 (26). Generation of Florida-based *D. simulans* line Lara-10 described in Zhao et al., 2014 (5). Generation of Zuma Beach, CA *D. simulans* line SZ116 described in Signore et al. 2018 (27) We also used the *D. yakuba* reference sequence strain, Tai18E2 (28).

With our greater sample size, we identified more unannotated genes expressed in AG tissue than found previously. A total of 90 genes were expressed exclusively within *D. melanogaster* lines (S3 table), and another 123 genes were expressed in *D. melanogaster* as well as *D. simulans* and/or *D. yakuba* (S4 table). We refer to the set of unannotated genes only expressed within *D. melanogaster* as “*melanogaster* unannotated” (MU), and unannotated genes present in *D. melano*gaster as well as *D. simulans* and/or *D. yakuba* as “multiple species unannotated” (MSU). We used a stringent filter for MU genes, excluding loci with any evidence of transcription in any *D. simulans* or *D. yakuba* genotype. Both MU and MSU categories include candidate *de novo*-evolved genes, since no evidence is present in our data that any species outside the *melanogaster* subgroup have orthologous sequences undergoing transcription. Since MU genes are restricted to a single species, their origin is likely very recent, and has likely occurred after the *D. melanogaster* – *D. simulans* split (Figure 1B). However, given the difficulty of sampling rare alleles, this category might contain some older genes as well (see next section). The MSU class is likely enriched for relatively young genes as well. However, since their origin is deeper in the past, there is an increased possibility that some of these are older genes that have undergone extensive sequence divergence, such that sequence homology can no longer be detected, or that individuals expressing these genes in more distantly related species were not sampled in our study.

**Figure 1:**
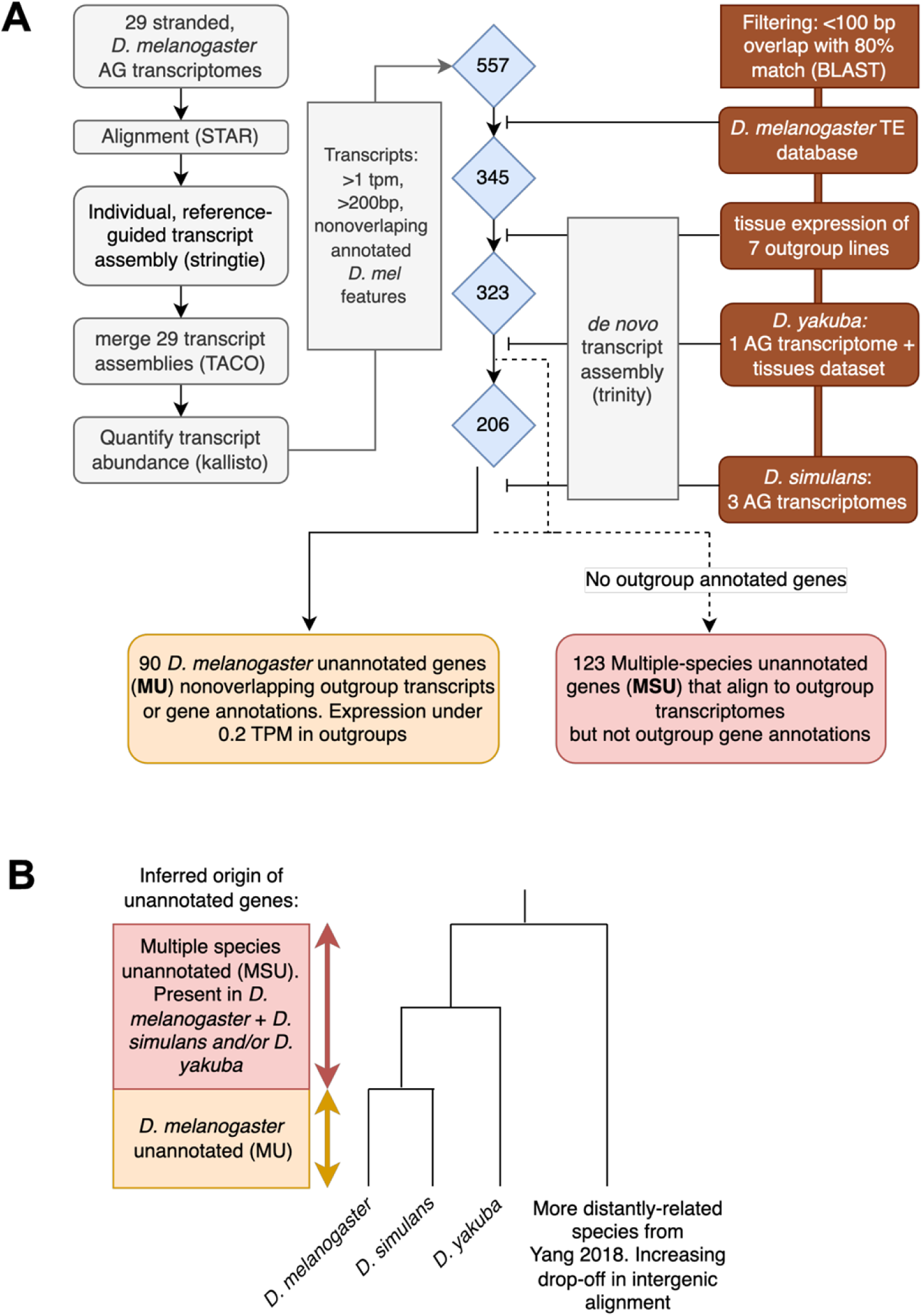
Candidate *de novo* gene discovery and species comparisons. (A) Workflow of candidate *de novo* gene discovery. Left column shows steps for identifying genotype-specific transcripts expressed in AGs. In brief, individually aligned transcriptomes were also individually assembled and then merged, across 29 DGRP genotypes, into one unified set. This assembly was appended to the *D. melanogaster* reference genome annotation 6.41 and then used to re-quantify transcript abundance across all individual *D. melanogaster* lines. Three preliminary filters were applied to the set of unannotated transcripts: they shared no overlapping bases with any annotated transcripts from *D. melanogaster* genome v6.41, were expressed at least 1 TPM in at least one *D. melanogaster* DGRP line and were at least 200 bp long. Next, we sequentially filtered out transcripts that showed >100 bp alignment and 80% sequence identity with any of the following expressed features, arguing against *de novo* origin in *D. melanogaster* (red boxes on right): transposable element sequences from the *D. melanogaster* v6.41 annotation; reads mapping to unannotated transcripts expressed in the 7 outgroup species sequenced in Yang et al. 2018 (52) (see table S2 for list of species and tissues in this dataset); *D. yakuba* r1.3 annotated transcripts, unannotated transcripts from Yang et al 2018, and unannotated transcripts from our dataset; *D. simulans* r2.02 annotated transcripts and unannotated transcripts from our dataset. For outgroup species, we constructed *de novo* transcriptome assemblies using Trinity (v2.11) (60) for each separate genotype or tissue expression sample (see S1 table for list of genotypes). After all filtering steps, we identified 90 *D. melanogaster* specific unannotated genes, and 123 genes that aligned to *D. melanogaster* plus *D. simulans* and*/*or *D. yakuba* transcriptomes but did not align to annotated genes in any outgroup species. (B) Phylogeny showing estimated age-range of unannotated genes identified in this study. Node for *D melanogaster* / *D. simulans* has been estimated ∼1.3 Ma; node for *D. melanogaster/D. yakuba* (*melanogaster* species subgroup) estimated ∼3.5 Ma (61).

Next, we examined the distribution of unannotated genes within *D. melanogaster*. We found that unannotated genes were typically expressed in very few *D. melanogaster* lines (Figure 2A-C), which supported results from previous studies (5,14,18). Across all MU genes, the average proportion of lines expressing each gene (at >1 TPM) was just 14%. This proportion was significantly higher for MSU genes, at 24.4%. This difference between age classes could reflect a gradual increase in the mean fitness of *de novo* gene sequences, with rare, maladaptive genes being purged over time. Yet the overall pattern was similar in both, with many genes expressed in just 1/29 lines. Only 2/90 MU genes and 1/123 MSU genes were fixed in *D. melanogaster* (present in all *D. melanogaster* genotypes). Together, these results are consistent with previous studies that suggest a high rate of turnover for *de novo* genes (29,30) and show that many of the alleles for both “younger” and “older” *de novo* genes may be close to being lost. One difference between this and previous studies is the increased sample of rarer genes, which provides better resolution for this overall trend.

**Figure 2:**
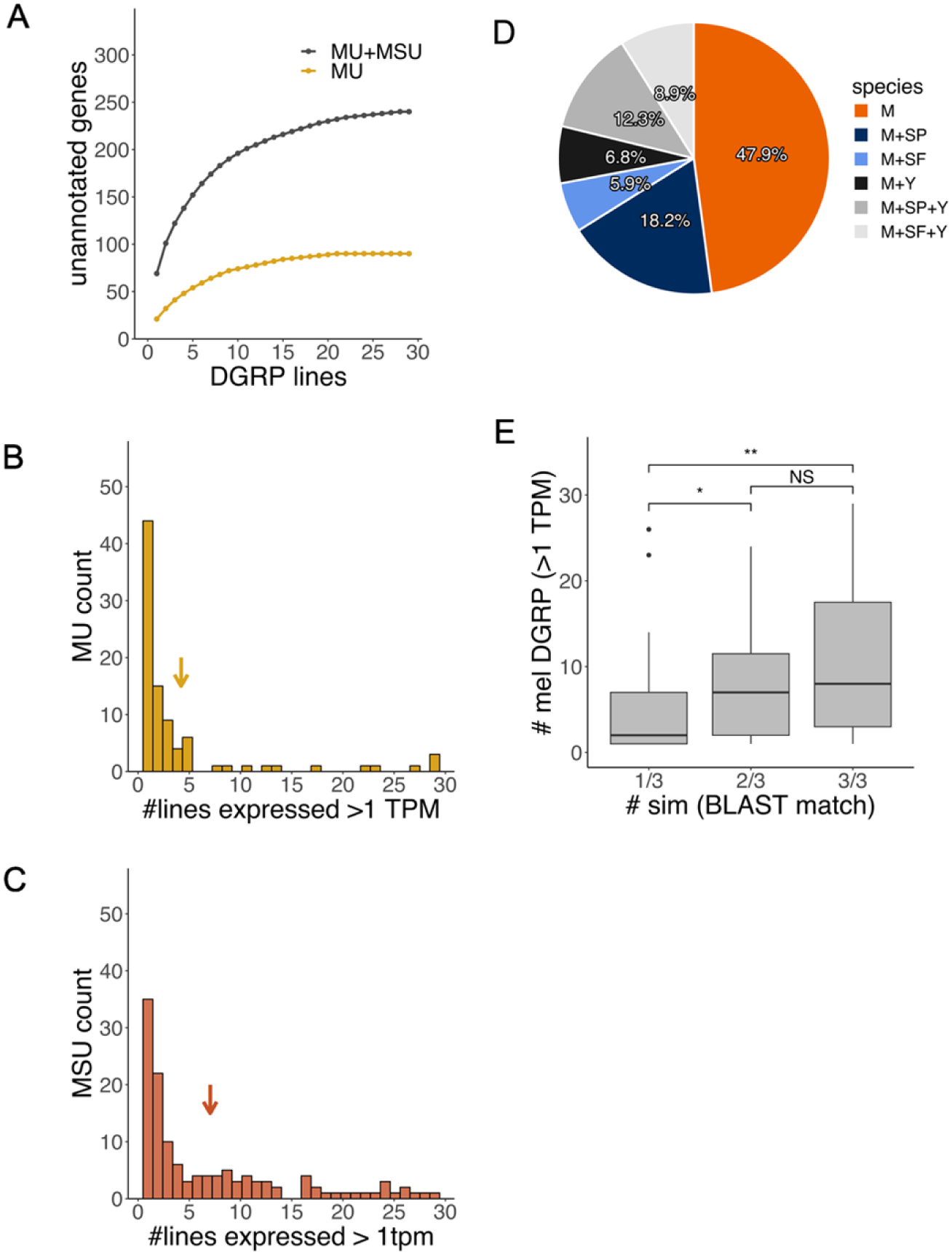
Most unannotated genes are at low frequency in the population. (A) Rarefaction plot depicting unannotated genes discovered per new genotype. Lines with more unique *de novo* genes are plotted first. Only 2/90 MU genes and 1/123 MSU genes were present in all 29 DGRP lines. (B, C) Comparison of MU (B) and MSU gene frequencies in the *DGRP*. Arrows indicate mean values. Expressed alleles at cutoff >1 TPM are at significantly higher frequency for MSU genes than for MU genes (Wilcoxon rank sum test with continuity correction; p<0.001). (D) Expression distribution of MSU genes in *D. simulans* and *D. yakuba*. Presence in outgroup genotype was established through transcript overlap of candidate *D. melanogaster de novo* genes with *de novo* transcript assemblies of outgroup AG transcriptomes. M: *D. melanogaster*; Y: *D. yakuba*, SP= *D. simulans* polymorphic (1/3 or 2/3 lines), SF= *D. simulans* fixed (3/3 lines). (E) Unannotated transcripts expressed in *D. simulans* genotypes are more likely to be rare in DGRP when they are also rare in *D. simulans* (Wilcoxon rank sum test with continuity correction; *p<0.05, **=p<0.01).

The two non-DGRP samples (iso-1 and ed10) also expressed the MSU genes that were fixed within the DGRP population. In relation to the DGRP genotypes, the African line (ed10) was on the high end for both number of expressed MSU genes and the total abundance of their transcripts (Figure S2). A high abundance of *de novo* genes expressed in an African sample has been reported previously (14). This suggests that many MSU genes originated prior to the spread of *D. melanogaster* from Africa and may be further evidence of a lack of expression divergence between North American and African in the AGs compared to other tissues (31).

### Expression of unannotated genes across species

The rarity of unannotated genes in *D. melanogaster* poses a potential difficulty for phylostratigraphic determination of gene origin. If a low proportion of individuals express a gene, it is unlikely that this gene will be found within a small, random sample of genotypes, leading to inaccurate origin estimates. For instance, if a gene exists in two species, but the rare genotypes expressing that gene were by chance sampled in only one of them, this gene may be incorrectly inferred to have originated after the two species diverged. We therefore investigated to what extent MSU genes are difficult to sample in *D. simulans* and *D. yakuba*, close relatives of our focal species *D. melanogaster*. If MSU genes are found at high frequencies in the outgroup species, it would strengthen our assignment of MU genes as *D. melanogaster*-specific and suggest that additional sampling would not affect estimates of their origin time.

Most MSU genes were not expressed in all individuals within our limited sample of *D. simulans* lines (Figure 2D). In addition, MSU genes expressed in fewer *D. melanogaster* lines were likely to be expressed in fewer *D. simulans* lines, as well (Figure 2E). We used permissive cutoffs for inferring the presence of *D. melanogaster* unannotated genes within *D. simulans,* requiring only a partial BLAST match to the *D. melanogaster* transcript with no minimum TPM cutoff. With this permissive cutoff, we found that 37/108 of *D. simulans*-expressed MSU genes were present in all 3 sampled lines. The genes that were found in 3/3 *D. simulans* genotypes were expressed in a significantly greater proportion of *D. melanogaster* lines than the genes present in only 1/3 *D. simulans* lines (Figure 2E). The correlation of expression frequencies between species suggests that rare, hard-to-sample genes are more likely to be miscategorized as species-specific, and that additional sampling of outgroup species may increase the accuracy of their inferred origin times.

We investigated whether the proportion of individuals expressing unannotated genes was related to their transcript abundance. The maximum TPM value measured, across lines, was significantly correlated with the number of DGRP lines expressing MU and MSU genes at a cutoff of >1 TPM (Figure 3A-B). This correlation between expression level and the frequency of expressed alleles in the population could indicate that expression variance affects unannotated gene calls. For instance, if mean expression is near the detection threshold, normal variation – either biological or technical – could result in transcripts being detected only in a fraction of individuals.

**Figure 3:**
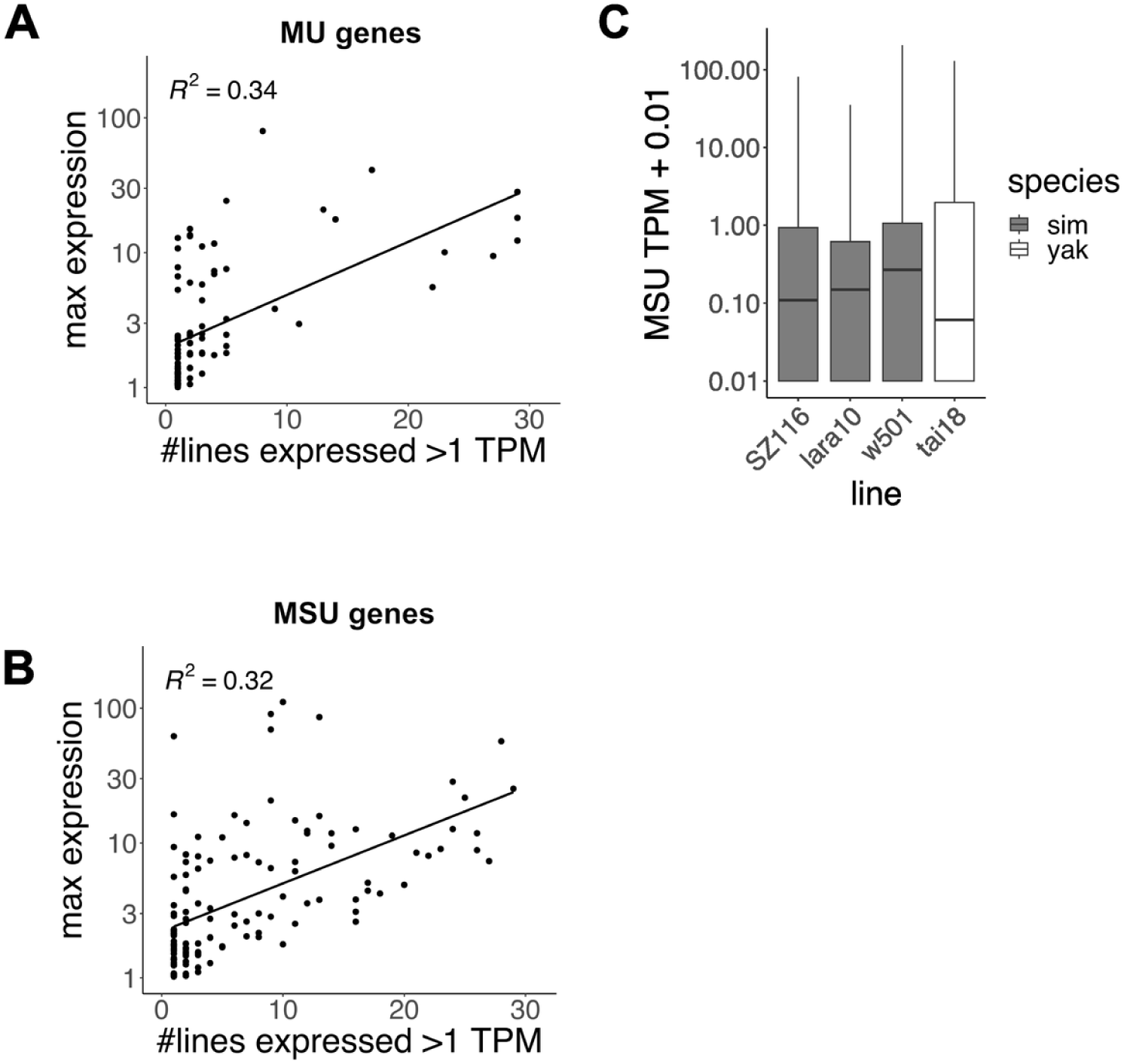
Unannotated genes with higher expression abundance are sampled in more genotypes. (A, B) Correlation between maximum expression value measured per unannotated gene and number of lines expressing that gene at a cutoff of 1 TPM. Both MU (A) and MSU (B) genes show significant correlation (Pearson’s product-moment correlation; p<0.0001). (C) Distributions of expression values of MSU genes in outgroup lines.

Other evidence suggests a continuum of unannotated expression in the genome, between multiple species, that may affect how unannotated genes are sampled. Different libraries of *D. simulans* line w501 and *D. yakuba* line tai18, sequenced by different lab groups, did not contain exactly the same transcript complement following *de novo* transcript assembly – though libraries from the previous study also included ejaculatory duct tissue (18). Expression of many MSU genes was very low in these species, thus they were plausibly near the threshold for transcript assembly (Figure 3C). And while the *D. melanogaster* specific (MU) category contained no genes that could be assembled in *D. yakuba* or *D. simulans*, many of these genes had a few reads that mapped to orthologous regions. 29/90 MU genes had very low, but still detectable expression (<0.2 TPM) in *D. simulans* and/or *D. yakuba*. Together, these results illustrate that unannotated gene regions do not always fit neatly into a binary “on” or “off” state for all individuals of a species. What exactly constitutes a species-specific *de novo* gene may be sensitive to the depth of sampling in outgroup species as well as to filtration parameters in both focal and outgroup species.

### Properties of de novo candidates and other unannotated transcripts

We expect that (1) new genes that are more advantageous will persist longer than less advantageous genes, (2) low frequency transcripts are more likely to be deleterious, and (3) genes present in multiple species are older than genes that are restricted to just one. Thus, differences between species-restricted and older genes could highlight the features that promote the persistence of *de novo* genes. In principle, such differences could reflect either differential retention of some types of *de novo* genes over others, or a gradual refinement of *de novo* genes by selection. Therefore, we compared various genomic properties of MU and MSU genes to explore how the characteristics of *de novo* genes may change over time.

First, we examined the tissue enrichment of MU and MSU genes. We used a dataset of tissue-specific RNA-seq from six different *Drosophila melanogaster* tissues (32) in order to calculate τ values of expressed genes (>1 TPM in any tissue). That study included two DGRP (R399 and R517) lines in common with ours. We found that a majority of AG-expressed unannotated genes were expressed at the highest abundance in AGs and exhibited tissue-restricted expression (Figure 4A). The proportion of AG-enriched genes was greater in the MSU category of unannotated transcripts, though this effect was not significant (p=0.211, Fisher’s exact test). Consistent with male-biased expression, the tissue with the second-highest median expression was the testis (Figure 4B). Together, these results suggest that the expression of a majority of AG-expressed unannotated genes from transcription factor activity that is enriched within the male reproductive tract, though the limited sample of MU genes expressed within lines R399 and R517 may be limiting our power to ascertain whether this changes significantly over evolutionary time.

**Figure 4:**
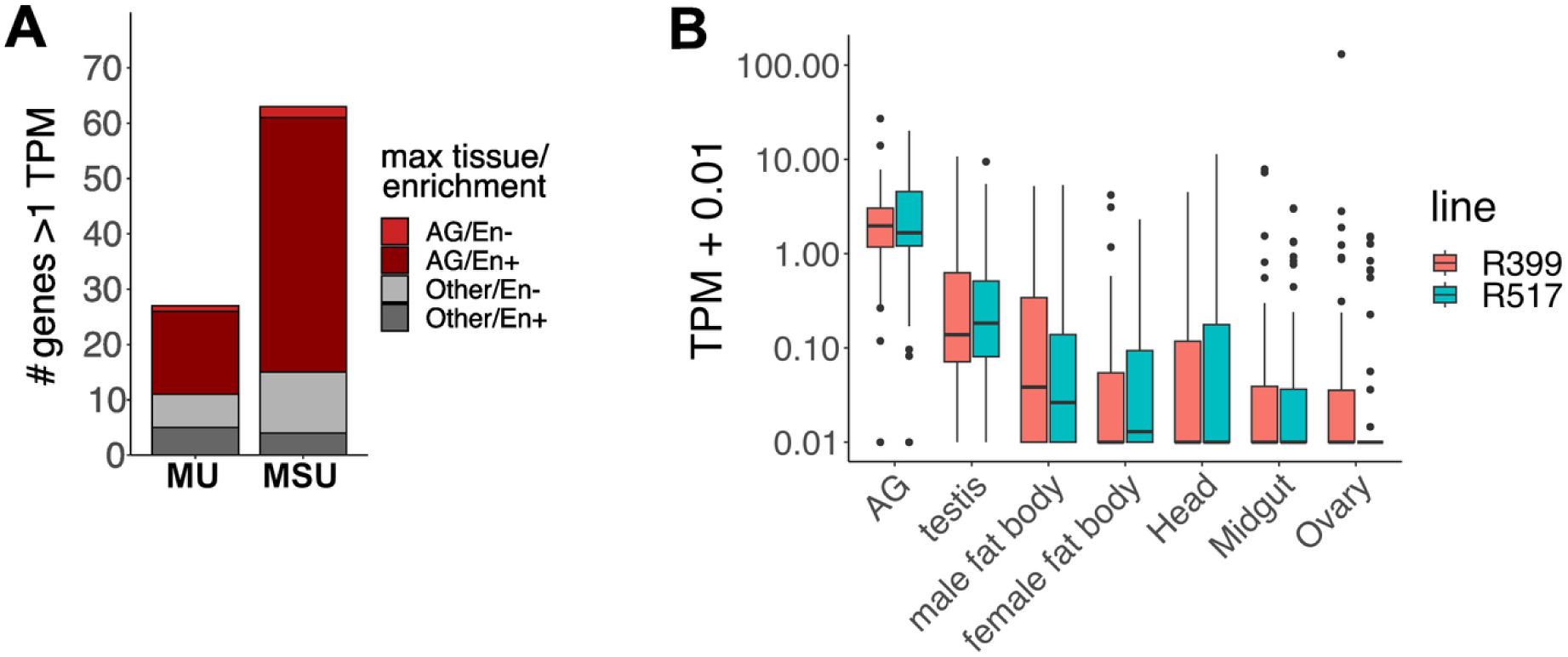
Many unannotated transcripts exhibit tissue-restricted expression. (A) Number of genes that are expressed at the highest level in AGs compared to other tissues, and exhibit AG enriched (En+) vs more widespread (En-) expression activity across tissues (cutoff for enriched expression: τ >0.9). No significant difference was found between MU and MSU counts for the four different categories shown (Fisher’s exact test; p=0.211). (B) Expression of unannotated transcripts (MU and MSU together) across tissues in two *D. melanogaster* lines.

Next, we explored several structural features of unannotated genes. Since a majority of both MU and MSU genes appear to have expression domains that are restricted to AG tissue, we also investigated how they compare to annotated, AG-restricted genes present in the *melanogaster* subgroup species (Figure S4). Both MU and MSU genes were expressed several orders of magnitude lower than typical AG-restricted, annotated genes (Figure 5A). All types of AG enriched genes (both annotated and unannotated) contained fewer exons than non-AG enriched genes (Figure 5B; pairwise Wilcoxon rank sum test with holm correction; MU vs AN p<0.001). Candidate *de novo* genes generally lack long predicted protein-coding ORFs (table S5), confirming the results from the previous study (18). Of MU and MSU genes containing ORFs (25 amino acid minimum length), there was no significant difference in the longest ORF per gene between classes (Figure S5). The proportion of MU genes that contained ORFs with predicted signal sequences was slightly lower: 4/90 MU genes vs 12/123 MSU. It is not clear how predictive the metrics of expression abundance or ORF presence are of gene function or “importance”. However, these results are consistent with a low impact of unannotated genes on the AG transcriptome, and likewise are consistent with many studies suggesting low levels of transcription and a lack of long ORFs in *de novo*/orphan genes (29,30,33–35).

**Figure 5:**
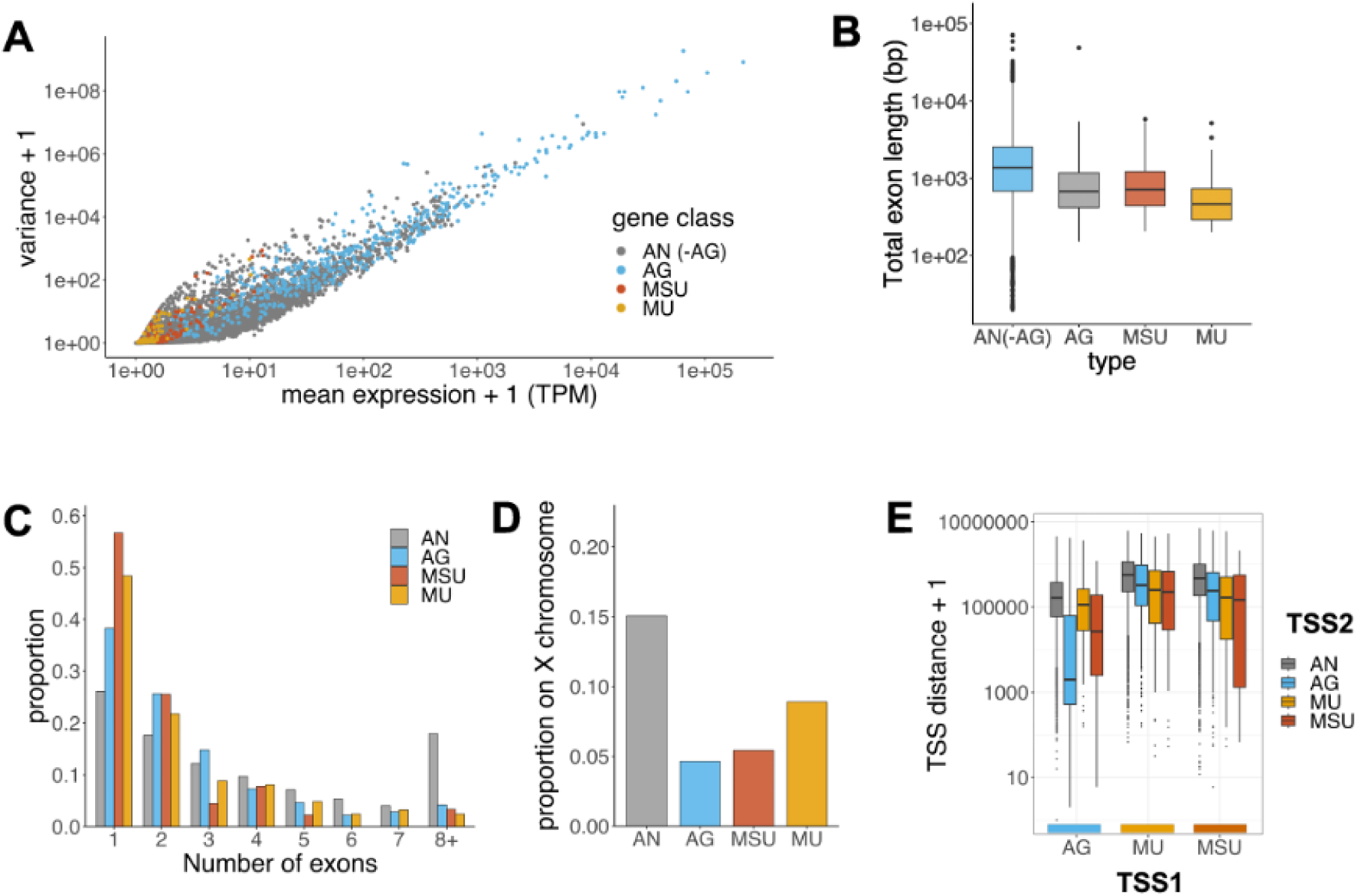
Annotated AG genes share several properties with unannotated genes, particularly with those that are older. (A) Most candidate *de novo* transcripts exhibit low mean expression with high relative expression variance compared to annotated genes. Abbreviations: AN (-AG): all annotated genes excluding the set of AG enriched genes; AG: Accessory gland enriched annotated genes; MSU: unannotated, expressed in *simulans* or *yakuba*; and MU: unannotated, restricted to *D. melanogaster*. (B) Gene length distribution, with MU genes being the shortest of all types (pairwise Wilcoxon rank sum test with Holm adjustment; p<0.001 for all contrasts). Non-AG-enriched, annotated genes are longer than other gene classes (p<0.001 for all contrasts), but annotated AG-specific genes are not significantly longer than ancestrally expressed unannotated (MSU) transcripts (p=0.57). (C) Exon numbers of full-length transcripts. MSU genes have fewer exons than MSU genes and annotated genes (Pairwise Wilcoxon rank sum test with Holm adjustment). Annotated, AG-specific genes do not contain significantly more exons than unannotated ancestrally expressed genes. Arithmetic means, per gene class, of the number of exons were: AN = 4.84, AG= 2.68, MSU = 2.67, and MU= 1.91 (D) MU genes are not as strongly depleted on the X chromosome as other AG-enriched gene classes. Though at a lower frequency, *de novo* genes were not significantly depleted on the X compared to annotated genes with less restricted tissue expression patterns after correcting for multiple comparisons (pairwise Fisher exact tests with Holm correction for multiple comparisons; MU vs annotated p=0.484; MSU vs annotated p=0.011; AG vs other annotated p<0.001). (E). Distance to the closest TSS, by gene class. Orientation of either transcript was not factored into this calculation.

There were other properties that distinguished the (presumably youngest) MU genes from other gene types. First, MU genes were significantly shorter (Figure 5C). Second, MU genes were not significantly depleted for the X chromosome, whereas MSU and AG genes were (Figure 5D; pairwise Fisher exact tests with Holm correction for multiple comparisons: MU vs annotated p=0.19; MSU vs annotated p=0.011; AG vs other annotated p<0.001). Third, nearest-neighbor analysis indicates that MU genes are located further away from annotated AG genes; in contrast, annotated AG genes are located closer to other annotated AG genes and MSU genes (Figure 5E). Together, these differences between MU and MSU genes could indicate the effects of directional selection on young *de novo* genes, where those that fit the “profile” of older AG genes may be more likely to persist over time.

### Bidirectional transcription of AG-expressed genes

We found 8/90 MU genes and 14/123 MSU genes that occurred within 1kb of another gene of the same class. This was an interesting finding, since multiple gene origin events within the same region could indicate a single genetic change precipitating the evolution of multiple *de novo* genes. To assess this, and to further determine whether *de novo* genes are reusing existing regulatory regions, we looked at a possible effect of bidirectional promoters as a mechanism of *de novo* gene origin (22). For this analysis, we used a previous definition of two –/+ oriented genes within 1 kb (36).

First, we noted that the proportion of genes fulfilling this cutoff for bidirectional transcription was a minority in any class of genes (Figure 6A). We compared the observed rate of cooccurrences to the distribution expected when cooccurrences are predicted based solely on the fraction of bidirectionally transcribed genes from each class. The rate of within-class cooccurrences was disproportionately high, indicating that bidirectional gene pairs are more likely to exhibit similar patterns of tissue-restricted expression (Figure 6B). Previous studies suggest that local “pockets” of AG-specific transcription exist in the genome (37,38). Consistent with these studies, we found a particularly biased distribution of AG-enriched annotated genes in pairs with other AG-enriched annotated genes: 31.8% of AG-enriched annotated genes are in bidirectional pairs with another AG-enriched gene, despite AG-enriched annotated genes constituting 2.5% of the genes examined.

**Figure 6:**
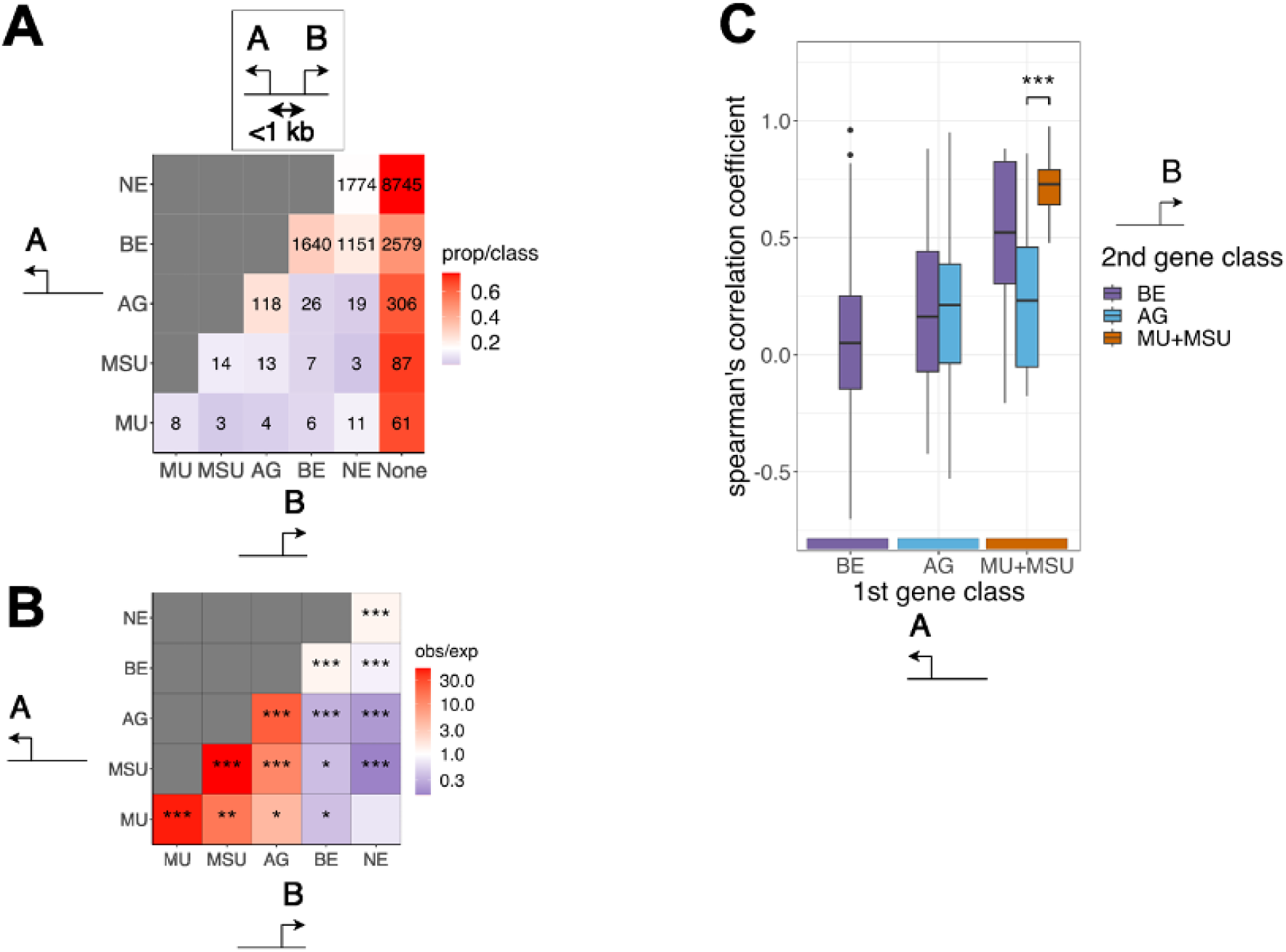
Genomic position of the youngest *de novo* genes does not support their origin via reuse of existing regulatory elements. (A) Frequencies of each gene class in divergently transcribed gene pairs. Each cell gives the proportion of genes in category A that occur in a bidirectional pair with a gene in category B (“none” indicates the proportion of singleton genes that are not part of bidirectional pairs). In this figure, each gene in a bidirectional pair is counted once as part of category A and once as part of category B (*e.g.,* eight *de novo* transcripts stemming from *de novo*-*de novo* bidirectional promoters, though only four promoters exist). MU: *D. melanogaster*-only unannotated genes; MSU: multiple-species unannotated genes; AG: accessory gland specific annotated genes; BE: subset of more broadly expressed annotated genes that are expressed, but not enriched within AGs; NE: annotated but not expressed in AGs. (B) Comparison between observed and expected counts of bidirectionally transcribed pairs, by gene class. Expected counts were calculated from the proportion of bidirectional genes in each class as a fraction of the total number of bidirectional genes. Asterisks indicate per-cell binomial test between observed and expected counts, with Holm correction for multiple comparisons (*p<0.05; **p<0.01; ***p<0.001). (C) Correlation of expression levels (Spearman’s ρ) between first and second gene in bidirectional promoter, by gene class. MU+MSU/AG pairs are significantly less correlated compared to pairs of unannotated genes in a bidirectional pair (p<0.001, ANOVA with Tukey HSD).

The number of bidirectional promoters containing either two MU genes or two MSU genes was approximately 50 times higher than expected (figure 6B). Although the sample size for these pairs was low, this observation could indicate a mechanism of gene origin where a single regulatory change spawns multiple new transcripts. When unannotated genes (particularly within the MSU class) occurred in a bidirectional pair with an annotated gene, the annotated gene was more likely to exhibit AG-restricted expression (Figure 6B). This suggests that tissue-restricted expression of unannotated genes (Figure 4A) may be influenced by the regulatory elements of existing AG-enriched annotated genes, particularly when a new gene evolves very close to an older gene.

Though the occurrence of bidirectionally transcribed *de novo* genes was low, it is possible that the enrichment of unannotated/AG-enriched bidirectional pairs reflects one of the mechanisms of *de novo* gene origin. Regulatory changes resulting in increased expression of the annotated gene could potentially cause concomitant transcription in the opposite direction, spawning a *de novo* gene in the process. In this scenario, we expect the expression levels of the annotated and the unannotated gene to be correlated. To test this hypothesis, we measured the extent of correlated expression of bidirectional gene pairs across the 29 DGRP lines. However, our results generally did not support this as a common mechanism of gene origin since unannotated genes in bidirectional pairs with AG-enriched genes exhibited the weakest correlations (Figure 6C). As such, the expression of *de novo* genes does not seem directly tied to existing regulatory activity, even if the occurrence of unannotated/AG-enriched gene pairs indicates that *de novo* genes can evolve in the permissive transcriptional environments near tissue-enriched, annotated genes.

Unannotated genes that were transcribed from bidirectional promoters did show strongly correlated expression in other instances. First, pairs of unannotated genes were the most strongly correlated of any class (Figure 6C; see figure S6 for example). This strongly correlated expression is consistent with a shared regulatory origin for these gene pairs. Second, a few unannotated/annotated gene pairs exhibited highly correlated expression, particularly when the annotated gene did not exhibit AG-enriched expression. For instance, the annotated gene CG45237 was polymorphic for expression in AGs, and thus not a part of the AG-enriched gene category. The lines that expressed CG45237 in AGs were more likely to express a nearby MU gene in the opposite direction (Figure S7).

## Discussion

In this study, we characterized recently evolved *de novo* genes expressed in *Drosophila melanogaster* male accessory glands. By sampling many more genotypes than previous studies, we were able to provide a better estimate of the frequency of *de novo* genes in natural populations.

We demonstrate that most of them are quite rare and most are expressed at very low levels. An improved, though still limited sampling of outgroup species revealed that some *de novo* genes may persist in populations for a long time without becoming fixed or even reaching high frequencies. These genes may be present in a polymorphic state in two or more related species, and therefore some *de novo* genes may be older than they appear to be based on shallower sampling. We found several ways in which the locations of *de novo* genes change as they age, suggesting several factors that may increase the likelihood of their persistence. The rarity of close association of *de novo* genes with either old AG-specific genes or bidirectional promoters suggests that most *de novo* genes are not simply repurposing existing regulatory regions that exist close to existing tissue-specific genes.

### Sampling and estimating the age of de novo genes

Our larger sample of genotypes greatly increased *de novo* gene discovery. We found 90 *de novo* genes using 29 DGRP lines, compared to 49 intergenic *de novo* genes from six lines in a previous study of the same tissue (18). Though a few of these additional candidates were due to relaxing gene distance cutoffs, our results are consistent with the prevalence of rare *de novo* alleles that were identified only by sequencing more genotypes. Even so, our study explored only one tissue under one set of environmental conditions, suggesting that many young, polymorphic *de novo* genes remain to be identified in *D. melanogaster*.

There is a complex relationship between genetic variation that leads to the emergence of polymorphic *de novo* genes and our ability to detect these genes in transcriptome samples. Previous allelic imbalance experiments found the expression of *de novo* genes to be partially explained, in most cases, by *cis*-regulatory variation, suggesting a local genetic basis for their expression (5,18). However, genetic variation can affect expression variance among genetically identical cells (“expression noise”) as well as mean expression level (39–41). In addition, differences in environmental conditions, random variation in RNA isolation and library preparation, and other technical factors can lead to a range of TPM estimates around the true mean. Since most *de novo* genes have low mean expression, increasing the number of samples increases the chance of sampling a replicate with TPM above the detection threshold. This will be true regardless of which genotypes are sampled, and even if genetic variation does not affect the mean expression level. Still, genes with higher mean expression will tend to have a higher proportion of replicates with TPM above the detection threshold, which could explain the observed correlation between expression level and estimated frequency of MSU genes (Figure 2E). However, segregation of ancestral polymorphisms or similar selective forces operating in multiple species may also contribute to this pattern.

A key conclusion from our work is that sequencing more genotypes reveals previously underappreciated uncertainty about the age and phylogenetic distribution of *de novo* genes. With a larger sample of genotypes, we can see that most *de novo* genes are expressed in very few lines. This is consistent with our understanding of *de novo* genes – they undergo rapid evolutionary turnover (14,30,42) and few reach high frequencies in populations. Yet as a technical consideration, rare occurrences are more challenging to sample. If it is difficult to sample individuals where a gene is expressed in one species, how certain can one be that this gene is entirely absent in another? For instance, a *de novo* gene truly expressed in 1/29 (3.4%) *D. melanogaster* individuals could easily go unsampled in our study. Given that the proportion of lines expressing a gene is correlated between species, what if this *de novo* gene is expressed in 3.4% of the *D. simulans* population as well? If so, our smaller sample of *D. simulans* genotypes is almost certainly leading us to overestimate the proportion of *de novo* genes that are exclusive to *D. melanogaster*, and thus underestimate the typical age of polymorphic *de novo* genes. Future studies would benefit from a more extensive sample of genotypes from multiple closely related species.

With more extensive intraspecific sampling, we can consider a range of alternative scenarios of gene evolution that can lead us to underestimate the age of polymorphic *de novo* genes or misinterpret other types of species-specific transcripts as *de novo* genes (Figure 7). The first two scenarios become more probable with greater divergence time between species: (1) fixed loss of a polymorphic gene in all individuals of the non-focal species and (2) sequence changes causing a failure to detect orthology. Our study calls attention to two more alternatives, stemming from difficulties of sampling gene expression: (3) ancestral polymorphic *de novo* genes may not be detected in outgroup species because of unsampled rare alleles and (4) low-abundance transcripts may not be sampled in the outgroups due to expression noise or technical variance, particularly in cases where the mean expression lies near the cutoff for detection. An underlying challenge with identifying *de novo* genes is that these alternative scenarios may not all be mutually exclusive.

**Figure 7:**
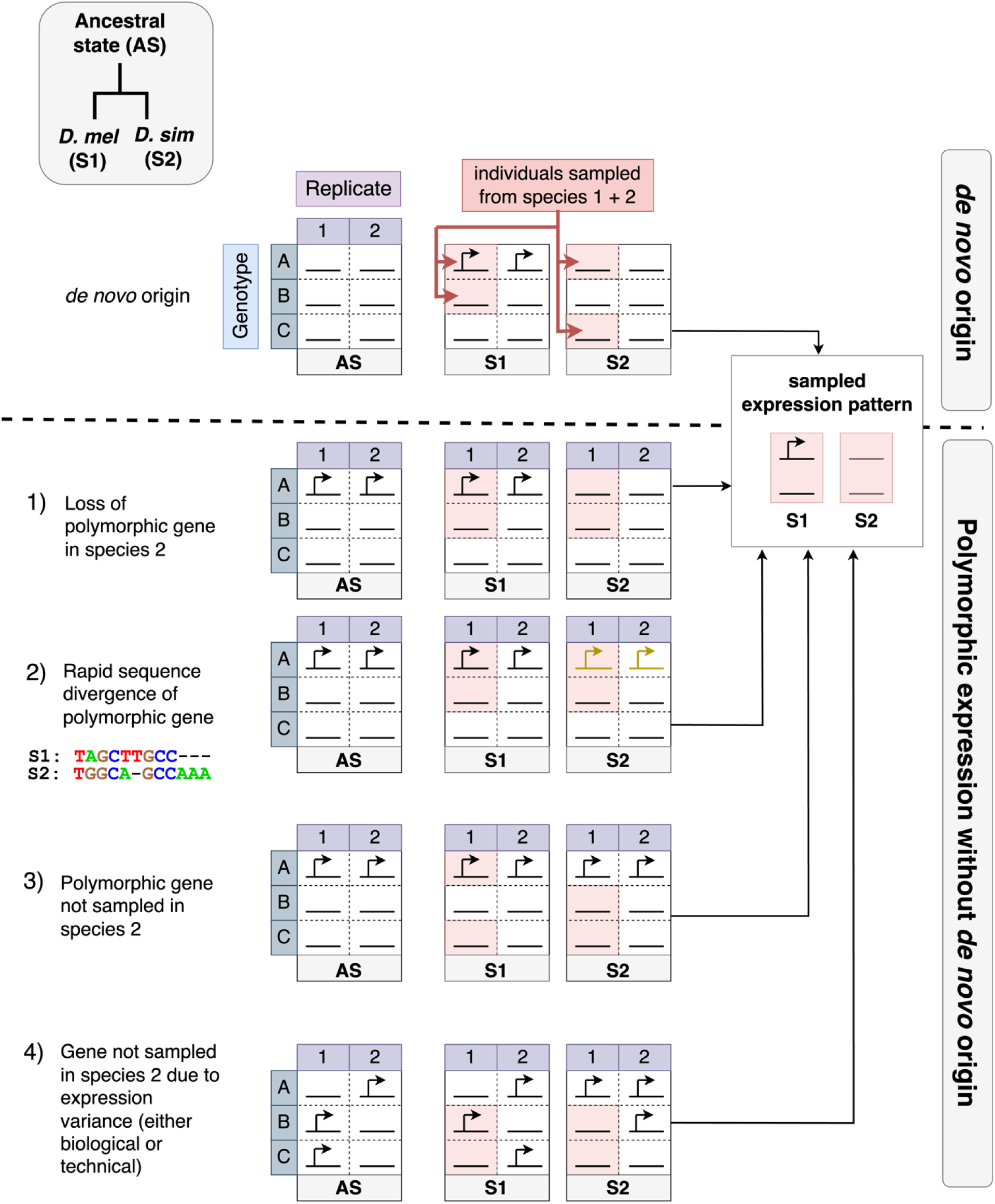
Four alternatives to *de novo* gene origin. The *de novo* origin of a gene is contrasted with four other scenarios leading to a pattern of polymorphic gene expression. In these examples, there are 3 genotypes (A-C) with 2 technical replicates each used to call expression. *De novo* origin: gene expression is a derived state exclusive to *D. melanogaster* and sampled only in one extant *D. melanogaster* line. Non *de novo* origin scenarios: 1) Fixed loss: ancestrally polymorphic gene is lost in all genotypes in the *D. simulans* lineage. 2) Sequence divergence: despite gene being present in some individuals of both *D. melanogaster* and *D. simulans,* failure to detect orthology leads to the appearance of a uniquely transcribed sequence. 3) Old Polymorphism: the ancestral state (the presence of a *de novo* gene in only some individuals of the population) is maintained in both *D. melanogaster* and *D. simulans* lineages. Since this gene may be rare, it is possible to miss evidence of its expression in one of the species. 4) Expression variance: highly variable expression of *de novo* gene within each genotype causes disagreement between replicates. Chance sampling leads to the appearance of gene loss in *D. simulans.* Note that in this case arrows indicate expression of gene, since all genotypes may be expressed under the right conditions.

In this study, we used low transcript abundance thresholds to improve the detection of young, rare *de novo* genes. In one sense, the degree of polymorphism observed here highlights the fallibility of relying on transcriptional evidence for assessing *de novo* gene origin, as suggested by others (43). Yet from another perspective, our results further showcase the widespread existence of polymorphic noncoding genes, even though how exactly to categorize low-abundance transcripts remains a gray area. Genes that both lack ORFs and are expressed at low abundance may reasonably be excluded as being too ambiguous in other studies focusing on *de novo* genes. However, the evolutionary transition from a nontranscribed to a transcribed state is key to *de novo* gene origin, with or without ORFs, though it may not be the most common limiting factor (44,45). For instance, a recent study in *Drosophila* found that the difference between active and non-active *de novo* gene alleles was most commonly transcriptional abundance alone, and not the composition of ORFs or structural differences within the gene (14). And yet, the question of “how much transcription is enough?” has no clear answer. The preadaptation hypothesis (46), for instance, suggests that *de novo* genes that evolve in transcriptionally “leaky” genomic regions are less likely to be extremely deleterious, since small amounts of transcription may still expose maladaptive alleles to selection. What exactly the meaningful distinctions are between “completely silent”, “ancestrally leaky transcription”, and “expressed enough to be categorized as a *de novo* gene” may be open to interpretation. To use *reductio ad absurdum*, if we set the expression threshold so low that any transcription at all is interpreted as a “gene”, no subsequent changes can serve as evidence for the evolution of a *de novo* gene at that location. Yet over a limited evolutionary span a majority of the genome may be transcribed (42). A discrete model for *de novo* gene origin may be useful in the absence of extensive intraspecific sampling. However, it does not easily allow for the possibility that some aspect of *de novo* gene evolution may involve stepwise increases in expression level that are more quantitative than qualitative. As we continue to explore the earliest steps in the evolution of *de novo* genes, this discrete model may give way to a more quantitative framework that focuses on the order and timing of multiple events that contribute to the emergence of increasingly robust transcripts.

### Positional and sequence composition clues about de novo gene regulation

We identified several systematic differences between younger and older polymorphic *de novo* gene candidates. On average, older genes are longer, less likely to be located on the X chromosome, and are located closer to other genes expressed in the same tissue. In all these respects, the older *de novo* genes are more similar to annotated AG-specific genes than are the younger ones. One possibility is that these differences reflect the work of directional selection. The observed locations of *de novo* genes are determined by two distinct forces: the probability that a new transcript will emerge in a given region, and the probability that this transcript will spread through the population. The initial origin of a *de novo* AG-expressed gene is more likely to occur far from any established AG-specific genes simply because most intergenic space is not near such genes. However, if the average *de novo* gene has increased fitness within a tissue-specific transcriptional environment, the few that do originate near older tissue-specific genes will be more likely to persist. In support of this hypothesis, prior studies in *Drosophila* found a correlation between lineage-specific gene age and tissue specificity (29,47). Similarly, *de novo* AG-expressed genes may be equally likely to emerge on any chromosome, but more likely to persist on the autosomes than on the X chromosome.

This study does not support the idea that *D. melanogaster de novo* genes commonly originate through reusing pre-existing regulatory elements near pre-existing genes. The youngest *de novo* genes typically exist further away from other annotated AG-specific genes (Figure 5E) and do not typically form close bidirectional associations within the promoter regions of annotated AG genes (Figure 6A-C). This was somewhat surprising, since these genes typically exhibit tissue-restricted expression (Figure 4A) and enhancers have been shown to facilitate *de novo* gene origin in mice (24). Such tissue-specific enhancers could provide a convenient “ready-made environment” that requires fewer changes to induce the transcription of a *de novo* gene. However, extremely close-by enhancers and bidirectional promoters may be a narrow subset of tissue-specific regulatory elements. These young *de novo* genes may be repurposing old enhancers that exist further within intergenic space, or conversely, they may be transcribed from entirely new enhancers. Four pairs of *de novo* genes stemmed from the same promoter region and tended to have highly correlated expression levels (Figure 6C). Transcription from enhancers in *Drosophila* is less “symmetrical” than in other systems, yet many enhancers do exhibit some bidirectional transcriptional activity (48). As such, these pairs of *de novo* genes could plausibly be enhancer RNAs stemming from new enhancers. Future studies of the genetic and epigenetic basis of *de novo* gene expression may evaluate the importance of these factors and provide alternative hypotheses for how *de novo* genes originate. Determining to what extent new genes coincide with new enhancer activity, for instance, could help identify the earliest causal changes leading to the origin of *de novo* genes.

## Methods

### RNA preparation

RNA libraries used for unannotated transcript discovery were constructed separately from the accessory glands (AG) of each of 29 lines of the Drosophila genetic resource panel (26). The DGRP consists of 205 sequenced isofemale lines derived from a single *D. melanogaster* population in Raleigh, NC. To assess the presence of unannotated transcripts in other *D. melanogaster* populations and to better establish patterns of outgroup expression, RNA was also collected from two non-DGRP *D. melanogaster* lines, two *D. simulans* lines, and one *D. yakuba* line (see Table 1 for lines used in this study, including previously published data). For each line, RNA was extracted from pooled accessory glands of 30 unmated, two day old males using TRIZOL (Invitrogen) followed by on column cleanup with DNAse digestion (Zymo). RNA libraries were prepared with Illumina Truseq stranded mRNA kit (Illumina), which uses polyT beads to capture polyadenylated RNA sequences. Libraries where sequenced using 150 bp paired-end reads on an Illumina Hiseq4000.

### Identification and quantification of de novo genes

To identify unannotated genes expressed in the accessory glands, 29 DGRP transcriptome sequences were aligned using STAR v2.6.1d (49) to the Dm6 genome assembly. A mean of 28,213,740 reads were mapped per sample, with a mean rate of 90.09% (S1 table). Reference-guided transcript assembly was performed for individual lines using stringtie version 1.3.4d (50). Next, assemblies from individual lines were merged into a unified file using TACO v0.7.3 (51) to create a preliminary set of transcripts expressed in accessory glands of DGRP lines. Two other *D. melanogaster* genotypes were sequenced from other populations (Table 1), but these were not included in the *de novo* gene assembly.

We then applied a series of filters to the assembled transcripts to identify the candidates most likely to have evolved recently in the *melanogaster* species subgroup. This procedure largely followed Cridland et al. (18), though, to examine the possible effect of pre-existing regulatory elements, we did not use any minimum distance cutoff between unannotated and annotated genes but excluded intronic sequences. See Figure 1 for description of filters used and overall workflow. We identified *Drosophila melanogaster* specific unannotated transcripts (MU genes) that were longer than 200 bp; were expressed >1 TPM in at least one DGRP line; aligned to at least 200bp of genomic sequence from *D. simulans* or *D. yakuba*; did not contain transposable element sequences; were located on chromosomes 2, 3, or X; and did not align to any transcripts (annotated or unannotated) from any of the 8 tested outgroup species (*D. yakuba, D. ananassae, D. pseudoobscura, D. persimilis, D. willistoni, D. mojavensis, D. virilis,* and *D. grimshawi*) in any of 8 tissues from a previous study (52). We also defined a second type of *melanogaster* subgroup specific transcripts (MSU genes) that passed the prior criteria, except these genes had greater than 50 bp alignment (at an 80% match) to any transcriptome sequence from either *D. yakuba* and/or *D. simulans*. Synteny analysis was performed for MU and MSU gene classes similar to Cridland et al. (18). For each candidate unannotated gene, we identified the first annotated gene immediately upstream and the first annotated gene immediately downstream. Next, we identified orthologs of these annotated genes using the Flybase 2021 ortholog database (Flybase; http://ftp.flybase.net/releases/FB2021_02/precomputed_files/orthologs/; downloaded May 12, 2021). We generated extended FASTA sequences of the candidate unannotated gene by extending the gene coordinates to include 5kb on either side, and we aligned this sequence to the region between the orthologous annotated genes + 10 kb on each side. For those genes that did not have programmatically confirmed synteny (due to, e.g., ambiguous ortholog calling or orthologs located on different scaffolds/chromosomes), we manually inspected corresponding genomic ranges on the UCSC genome browser using the latest assemblies and annotations (Prin_Dsim_3.1 Oct. 2021 fly *D. simulans* (w501 v3.1 2021) (GCF_016746395.2), Prin_Dyak_Tai18E2_2.1 Jul. 2021 fly *D. yakuba* (v2 2021) (GCF_016746365.2)).

We performed a more stringent final filtration step on MU genes by testing for transcription in the AG in the *D. simulans* and *D. yakuba* lines. Even if no corresponding transcripts could be assembled in *D. simulans* or *D. yakuba*, any genes with reads present at >0.2 TPM in either species were excluded from the final list of MU genes. Since these genes did not assemble into transcripts in either *D. simulans* or *D. yakuba*, they were not included in the MSU class either.

In total, these filtration steps yielded transcripts for 90 MU genes, which we provisionally consider as candidate *D. melanogaster*-specific *de novo* genes. They also produced a set of 123 MSU genes -unannotated ancestral transcripts whose origin likely predates the *D. melanogaster*/ *D. simulans* split.

To quantify the abundance of candidate *de novo* and annotated transcripts, we appended the unannotated sequences to the *D. melanogaster* reference annotation v6.41 and used kallisto (53) v0.46.2 to quantify transcript abundance in all DGRP lines, an African *D. melanogaster* line (ed10), and the *D. melanogaster* genome strain (iso-1). We used the tximport R package (54) to summarize transcript counts across genes and collect TPM measurements. To compare our calculated expression values to those of replicate samples from the same tissue but with slight differences in rearing and RNA library preparation methods, we also quantified transcript abundance in the data from Cridland et al. (18) and Zhang et al. (32) (see Table 1 for list of genotypes) using the same custom annotation file containing the appended unannotated transcripts (*i.e.,* we did not separately call *de novo* genes using these libraries).

### Coding potential

Coding potential of MU and MSU genes was assessed using Coding Potential Assessment Tool (CPAT) (55), using default parameters. For *Drosophila melanogaster*, the proprietary coding potential cutoff is set at 0.39, with a default minimum ORF length of 75 nucleotides. to assess candidates for signal sequences, protein translations were analyzed with SignalP 6.0 (56) using default settings.

### Tissue-specificity of unannotated genes

To quantify whether unannotated genes exhibit tissue restricted expression, we used RNA-seq data from a previous study within *D. melanogaster* (32). This dataset is comprised of six different tissues (head, male and female fat body, ovary, testis, midgut, and AGs) with two technical replicates per tissue, and contained data from the lines R399 and R517 which were also included in the AG transcriptomes generated for this study. To quantify expression in each separate tissue, we used the same transcriptome assembly generated from the AG-specific RNA seq data, and obtained TPM counts of transcripts as previously described using kallisto (53).

### Selection of AG-biased annotated genes

We selected *D. melanogaster* annotated genes (v6.41 *D. melanogaster* release) that demonstrated strong AG bias. This procedure was similar to Cridland et al. (18). We used the FlyAtlas2 data (57) to identify genes that showed the highest expression in the *D. melanogaster* AG and showed tissue specificity index (58) τ > 0.9. Next, genes with mean expression <1 TPM across DGRP lines were filtered out. We also removed genes that we could not verify originated prior to the *D. melanogaster – D. simulans* split. To accomplish this, we processed these annotated genes through the same BLAST pipeline used to identify *de novo* genes, except for the gene syntenty analysis. Fifteen annotated AG genes, all of them noncoding RNAs, which did not have any BLAST matches >100 bp to annotated genes or *de novo* transcriptome assemblies in the outgroup species were removed. In total, 452 AG-enriched *D. melanogaster* genes passed these criteria (Figure 2A).

### Analysis of bidirectional promoters

First, we identified all genes occurring in bidirectional pairs in the genome using a prior definition of two –/+ oriented genes within 1 kb (36). Next, we binned genes by annotation status, expression of unannotated genes (restricted to *D. melanogaster*, or expressed in other species as well), and tissue specificity (specific to AGs vs broadly expressed), and then counted cooccurrences of each gene class in bidirectional pairs. To obtain expected frequencies of cross-class bidirectional pairs, we calculated the joint probability of obtaining a bidirectional gene for each pair. For this analysis, each class was weighted by the number of bidirectional genes belonging to that class as a fraction of all genes within bidirectional pairs. Expected counts from pairs containing two different gene classes were doubled, as there are two different combinations that produce such pairs. We then sought to determine whether certain gene class pairs were more likely to be coregulated. To do this, we transformed transcript-level RNA counts using edgeR (59) to correct for sequencing depth differences between libraries and calculated spearman correlation coefficient between genes in divergently transcribed pairs. Due to the low counts of MU and MSU genes in bidirectional pairs, these two classes of genes were binned together for this analysis.

## Acknowledgements

We would like to thank Ben Hopkins, Olga Barmina, and Yuichi Fukutomi for providing helpful feedback on the manuscript; Sarah Signor and the Bloomington Stock Center for *Drosophila* strains; Li Zhao for initial guidance on the project; and Zachary Achilles, Ellie Lassotovitch, and Tammy Chan for assistance with fly care. The sequencing was carried out by the DNA Technologies and Expression Analysis Core at the UC Davis Genome Center, supported by NIH Shared Instrumentation Grant 1S10OD010786-01. This work was supported by NIH grant R35 GM122592 to A.K. and NIH grant R35 GM134930 to DJB.

## Data Availability

Accessory gland sequences available at https://www.ncbi.nlm.nih.gov/sra under BioProject accession number PRJNA1040452. Pipeline scripts and information have been uploaded to https://github.com/logankblair/AGdenovo.

## Supplemental captions

**Table S1: Mapping statistics of RNA seq.**

**Table S2: Species used for outgroup transcript screening**. Source of outgroup sequence data is from Yang et al. 2018 (52). Tissues for all species included: female abdomen without digestive or reproductive system, female digestive plus excretory system, female gonad, female reproductive system without gonad, female thorax without digestive system, female whole body, male abdomen without digestive or reproductive system, male digestive plus excretory system, male gonad, male head, male reproductive system without gonad, male thorax without digestive system, and male whole body.

**Table S3: Information about MU genes, including expression values.**

**Table S4: Information about MSU genes, including expression values.**

**Table S5: Predicted coding potential and signal sequences from MU and MSU transcripts**

**Figure S1: Synteny analysis of *D. melanogaster* unannotated genes.** Synteny confirmed through BLAST match of unannotated gene sequence to intergenic outgroup sequences anchored by next-nearest annotated genes (see methods).

**Figure S2: Number of genes expressed >1 TPM per line in *D. melanogaster***. Red bars indicate non-DGRP lines: ED-10 originates from Africa, and iso1 is the reference genome strain.

**Figure S3**: **Combined expression of all *de novo* genes (total TPM) per *D. melanogaster* line.** Red bars indicate non-DGRP lines: ED-10 originates from Africa, and iso1 is the reference genome strain.

**Figure S4: Tissue specificity of AG-expressed genes.** Data from our study showing mean AG expression (AG exp) of all annotated genes on the Y axis vs data from FlyAtlas 2 (Leader *et al.*, 2018) (57) showing tissue specificity (Sp) on the X axis, based on the tau index. Threshold for specificity (Sp) is set at 0.9, and expression is set at 1 TPM; horizontal dotted line indicates 2 TPM. Light blue points in the top right quadrant form the set of annotated, AG-enriched genes used in this study. Dark blue points correspond to genes that were expressed at the highest levels in AGs, compared to other tissues, in the Leader *et. al* 2018 study, but these genes were either expressed broadly in other tissues of their dataset, and/or exhibited low expression in our data (possibly related to the restricted age window and virgin status of males in this study).

**Figure S5: Distribution of ORF lengths is similar between MU and MSU genes.** No significant difference was detected in mean ORF length, minimum 25 amino acids, between MU (n=78) and MSU (n=115) genes (Welch two sample t test; p= 0.4696).

**Figure S6**: **Example of a DN-DN bidirectional gene pair.** IGV screenshot showing 5 expressed alleles and 5 non expressed alleles (in blue). “Unannotated transcripts” row shows transcripts corresponding to two separate genes (TU5897-TU5898 transcribed on Crick strand and TU5815 transcribed on Watson strand).

**Figure S7**: **Example of a DN-annotated gene bidirectional pair.** IGV screenshot showing 5 highest-expressed alleles of unannotated transcript TU5932 (expressed on Crick strand). Annotated gene is expressed on Watson strand in some, but not all, genotypes.

## Notes

### Competing Interest Statement

The authors have declared no competing interest.

